# Essential elements of radical pair magnetosensitivity in *Drosophila*

**DOI:** 10.1101/2021.10.29.466426

**Authors:** Adam A Bradlaugh, Giorgio Fedele, Anna L Munro, Celia Napier Hansen, Sanjai Patel, Charalambos P. Kyriacou, Alex R. Jones, Ezio Rosato, Richard A. Baines

## Abstract

Many animals use the Earth’s magnetic field (geoMF) for navigation^1^. The favored mechanism for magnetosensitivity involves a blue-light (BL) activated electron transfer reaction between flavin adenine dinucleotide (FAD) and a chain of tryptophan (Trp) residues within the photoreceptor protein, CRYPTOCHROME (CRY). The spin-state of the resultant radical pair (RP), and hence the concentration of CRY in its active state, is influenced by the geoMF^2^. The canonical CRY-centric radical pair mechanism (RPM) does not, however, explain many physiological and behavioural observations^2–8^. Here, using electrophysiology and behavioural analyses, we assay magnetic field (MF) responses at single neuron and organismal level. We show that the 52 C-terminal (CT) amino acids of CRY, which are missing the canonical FAD binding domain and Trp chain, are sufficient to facilitate magnetoreception. We also show that increasing intracellular FAD potentiates both BL-induced and MF-dependent effects on the activity mediated by the CT. Additionally, high levels of FAD alone are sufficient to cause BL neuronal sensitivity and, remarkably, potentiation of this response in the co-presence of a MF. These unexpected results reveal the essential components of a primary magnetoreceptor in flies, providing strong evidence that non-canonical *(i.e.,* non-CRY-dependent) RPs can elicit MF responses in cells.

## Introduction

The ability of species to navigate considerable distances has long intrigued the biological community^1^. One of several environmental cues to support these migrations is the geoMF. Additionally, several other behaviours respond reliably to MFs, at least under laboratory conditions, showing that the ability to sense and react to MFs is not limited to migrating animals^9^. However, the identity of the primary magnetoreceptor(s), the mechanism(s) that underlies its reported light dependence and how the magnetic signal is transduced remain unknown^10,11^. A favoured model posits a light-induced electron transfer reaction whereby RPs are formed, the spin-states of which are sensitive to MFs as small as the geoMF (~50 μT)^2^. This so-called RPM, canonically requires the flavoprotein CRY, which is best known for its role as a circadian BL-photoreceptor in flies and as a light-insensitive transcriptional regulator in the circadian clock of mammals^2,10^.

Absorption of BL by CRY-bound FAD initiates an electron transfer cascade along a conserved chain of Trp residues^2,12–14^. In *Drosophila* this forms a spin-correlated RP comprising the photoreduced FAD (FAD°^-^) and the terminal oxidised Trp (TrpH°^+^)^15^. The spin-state of the RP is initially polarised as a singlet (S, anti-parallel spins), which then rapidly oscillates between S and the triplet spin states (T, parallel spins). Transiently *(i.e.,* before the system relaxes to equilibrium), this inter-conversion can be sensitive to MF, which in turn can lead to downstream modifications in the biological activity of CRY, *via* conformational change^2^. In its activated state, the CRY C-terminal ‘tail’ of ~20 residues (CTT) becomes exposed, allowing interactions with signalling partners including PDZ-domain containing proteins^16–22^.

Although there is ample evidence consistent with CRY being both necessary and sufficient for light-dependent magnetosensitivity, there are a number of studies that support exceptions to this mechanism ^2–8^. In one of the most striking, Fedele and colleagues used a circadian behavioural assay in *Drosophila* to show that CRY-dependent light and magnetosensitivity could be rescued in CRY-null adult flies *via* expression of the 52 CT residues of CRY fused to GFP (GFP-CT) for stability^3^. Furthermore, CRYΔ, resulting from the deletion of the CTT of CRY, appeared largely insensitive to a MF, although BL sensitivity was maintained^3,4^.

The *Drosophila* CRY-CT lacks both the FAD-binding pocket and the chain of four Trp residues (W394, W342, W397, W420) presumed necessary for the canonical RPM^2,23–25^. Moreover, mutating these Trp residues, including W420F and W342F, at best attenuates, but does not abolish the magnetic functionality of CRY^3,4,26,27^. These results are inconsistent with current understanding of the RPM and question the identity of the magnetic-sensitive RP in the receptor. Proposed alternatives to a RP between FAD°^-^ and TrpH°^+^ include the formation of a RP between FAD°^-^/FADH° and O2°^-^ or another (unknown, Z°) radical. It is a matter of some contention whether these ‘unconventional’ RPs contribute to magnetoreception or even represent a primary sensor^11,28–30^.

Here we report the expression of a new transgene encoding Luc-CT (CRY-CT fused to luciferase), which lacks the canonical FAD-binding pocket and Trp chain and thus, is unable to support light-induced intramolecular electron transfer. Nevertheless, Luc-CT is sufficient to generate changes in BL and MF-dependent phenotypes in a whole organism circadian behavioural assay and in the electrophysiological activity of a model neuron, the larval aCC motoneuron. We show that the MF-responsiveness of Luc-CT is potentiated by increasing the intracellular concentration of free FAD, to the point where high levels of this flavin alone are capable, in the absence of Luc-CT, to support a MF response. Finally, we confirm by mutational analysis that the integrity of the CTT of CRY correlates with its ability to facilitate sensitivity to a MF. Overall, our results suggest that ‘sensing’ and ‘transducing’ MFs are separate properties that do not necessarily have to be carried out by the same molecule.

## Results and Discussion

### CRY-CT is sufficient to support magnetosensitivity

To validate our electrophysiological assay we expressed full-length *Drosophila* CRY (DmCry) in the aCC motoneuron; this supported a BL-induced increase in action potential (AP) firing by 1.7-fold and by 2.4-fold, in the co-presence of a MF (BL+MF,100 mT static, Fig.1A-B, p=0.005, see also Extended Data Fig.1a)^4^. Expression of CRY-CT (fused to GFP) in a *cry-null* background supports a MF-induced shortening of circadian period^3^. To eliminate the possibility that GFP might, like CRY, support intramolecular light-induced electron-transfer, we fused CRY-CT to Luciferase (Luc-CT) and maintained the flies in the absence of luciferin substrate. BL lengthened the free-running period of *tim-GAL4 > UAS-Luc-CT; cry^02^/cry^02^* flies compared to those in constant darkness (DD) (24.50 h *vs*. 23.75 respectively, p=0.019) revealing the BL light-sensitivity of CRY-CT. As with the GFP-CT construct^17^, Luc-CT exposure to a MF (300 μT, 3 Hz) was sufficient to shorten the free-running circadian period in the MF exposed but not the sham group (0 μT MF, Fig.1Ci, pre-exposure/post-exposure x sham/MF interaction F_1,377_=7.6, p=0.006, Extended Data Fig2.a-b). Notably, the Luc-CT fusion supports a marginal MF-mediated period shortening compared to sham even at 50 μT, which is geoMF strength (Extended data Fig.2f-j). Remarkably, expression of Luc-CT in the aCC neuron supported a BL-induced increase in AP firing (1.4-fold,), which was increased further in the co-presence of a MF (100 mT, Fig.1Cii, 2-fold, p=0.002, Extended data Fig.1b). In summary, these data collectively show that the CRY-CT alone is sufficient to support magnetosensitivity in both circadian and electrophysiological phenotypes. We predict that it does so through its well described interaction with the redox-sensitive K^+^channel β-subunit HYPERKINETIC (HK) ^30^.

**Figure 1.**
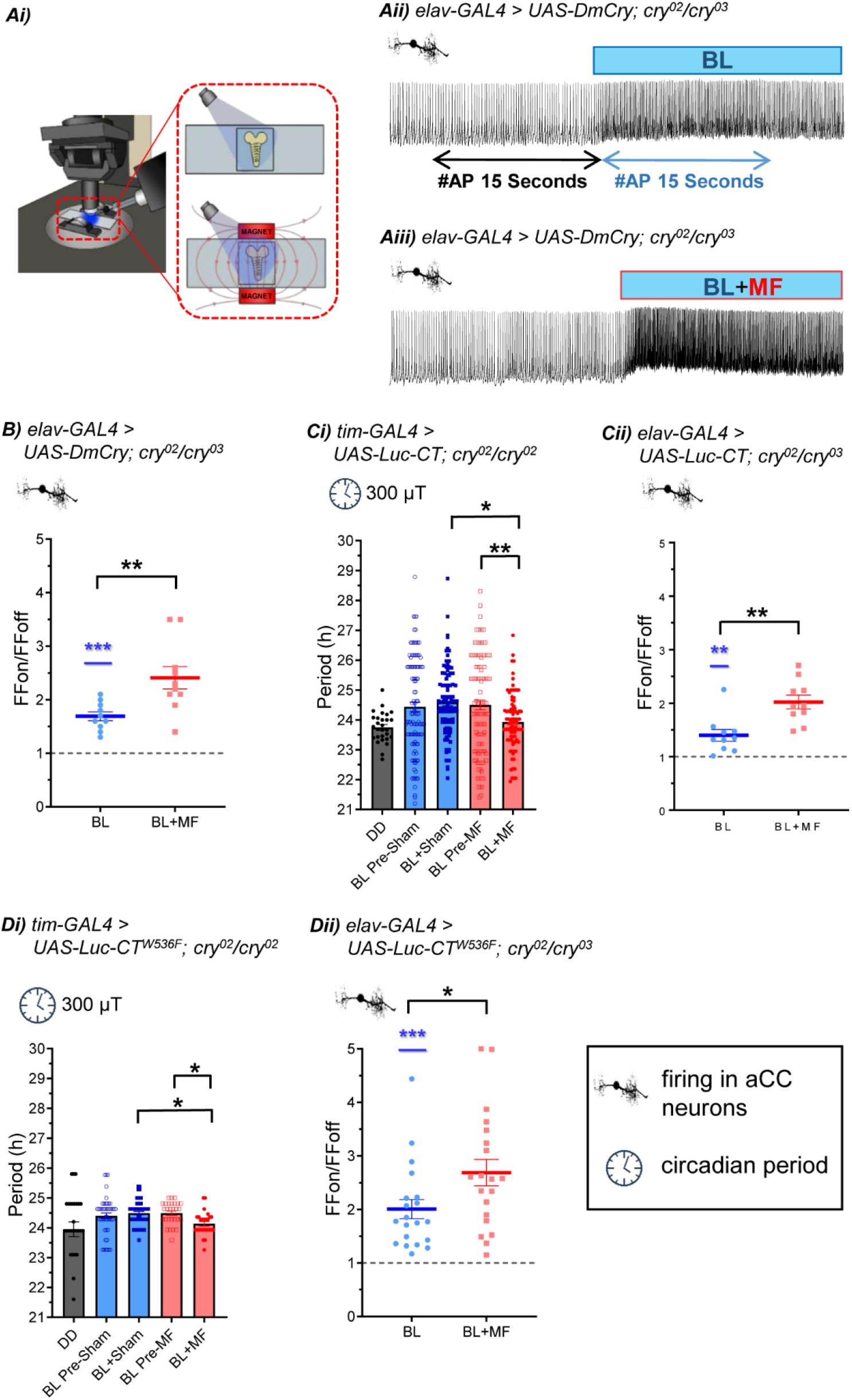
Luc-CT is sufficient to support magnetosensitivity. (Ai). Electrophysiological setup (permanent magnets shown in red) (Aii) BL-exposure of aCC neurons expressing DmCry increases action potential firing. (Aiii) Co-presence of MF (100 mT) potentiates this effect. Traces from different preparations. (B). Relative firing frequency of aCC expressing DmCry. BL increases firing 1.69-fold (t_(9)_=7.72, p=<0.0001, n=10, Firing-Frequency_on_/Firing-Frequency_off_). External MF (BL+MF, 100 mT) potentiates effect to 2.41-fold (BL *vs* BL+MF, t_(18)_=3.2, p=0.005, n=10, Extended Data Fig.1a). (Ci). *tim-GAL4>UAS-Luc-CT; cry^02^/cry ^02^* show period shortening under MF (Sham/MF x pre/post-exposure interaction (F_1,377_=7.6, p=0.006, 3-way ANOVA). MF-exposed flies show significantly shorter period than sham. Each of 4 replicates (Replicates, third factor in ANOVA) showed the same period shortening under MF (Extended data Fig.2a-b). (Cii). Luc-CT supports BL-induced firing (1.4-fold, t_(9)_=4.01, p=0.003 n=10, Extended Data Fig.1b) potentiated 2-fold in BL+MF (BL *vs* BL+MF, p=0.002, t_(df,18)_ =3.71, n=10). (Di). Luc-CT^W536F^ revealed significant period shortening on exposure to MF (significant pre/post-exposure x MF/sham interaction F_1,198_=5.1 p=0.025, 2-way ANOVA, *post-hoc* tests in Extended Data Fig.2c-e). (Dii). aCC expressing Luc-CT^W536F^ show 2-fold change in BL-induced firing (t_(19)_=6.06, p=<0.0001, n=20). Response to BL+MF is more variable, but significantly greater than BL alone (2.69-fold, 2-way ANOVA, Replicates as factor, p=0.03, F_(1,16)_=5.09, Extended Data Fig.1c). Raw data responses are shown in Extended Data Fig.1. Controls are reported in Extended Data Fig.3. For FFon/FFoff data, blue asterisks represent significance comparing before *vs*. during BL exposure (same cells, paired t-test); black asterisks represent comparisons of BL *vs.* BL+MF (different cells, unpaired t-test). * p=≤0.05, ** p=≤0.01, *** p=≤0.001.

### CRY CT is not a RP partner via Trp536

Although CRY-bound FAD may be dispensable, it is possible that FAD from another source in proximity could interact by forming a RP with the sole (non-canonical) Trp in the CRY-CT (Fig.3A). This alternative mechanism may explain why mutations of single Trp residues that constitute the Trp-tetrad are not significantly detrimental to CRY dependent magnetoreception^3,4,26,31^.

**Figure 2.**
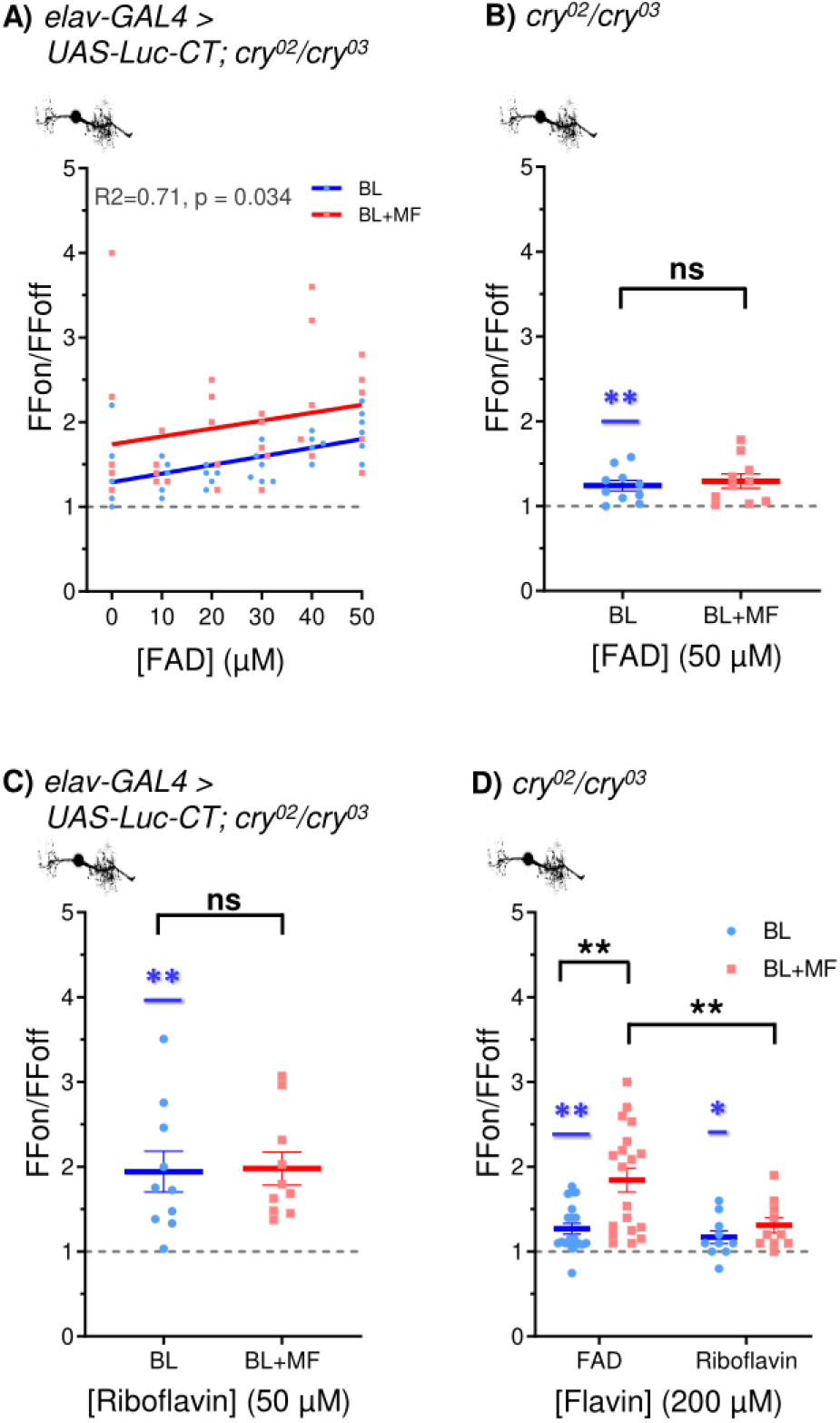
Free FAD potentiates the effect of Luc-CT and at high concentration supports magnetosensitivity alone. (A). Exposing aCC expressing Luc-CT to FAD (*via* the recording pipette) increases the response to BL (R^2^=0.71, F_(1,4)_=10.1, p=0.03). The co-presence of a MF (100 mT) potentiates the response (F_(1,9)_=9.06, p=0.015) by a consistent amount across FAD concentrations tested (n=5 except BL 30 μM and 50 μM [FAD] for which n=6). (B). Addition of FAD (50 μM) supports BL sensitivity (see also Extended Data Fig.4a), but not magnetosensitivity in the absence of Luc-CT (p=0.609, t_(18)_=0.521, n=10). (C). Addition of riboflavin (50 μM), supports the response to BL in aCC expressing Luc-CT, but not MF potentiation (t_(18)_=0.12, p=0.91, n=10). (D). Increased FAD (200 μM), in *cry^0^,* supports BL-induced change in firing (1.27-fold, t_(19)_=4.29, p=0.0004, n=20, Extended Data Fig.4c). The co-presence of a MF (n=19), significantly potentiates this level of FAD (1.84-fold, 2-way ANOVA (F_(1,55)_=3.51, p=0.066), Newman-Keuls *post-hoc* p=0.003). Riboflavin (200 μM) shows a similar BL effect (1.17-fold, t_(9)_=2.33, p=0.045, Extended Data Fig.4c) but no MF potentiation (1.31-fold, Newman-Keuls *post-hoc* p=0.67). Raw data is reported in Extended Data Fig.4. Blue asterisks represent significance values for aCC before *vs*. during BL exposure (paired t-test, same cells); black asterisks (or ns = not significant) represent comparisons of BL *vs* BL+MF (unpaired t-test, different cells). ns p=>0.05, * p=≤0.05, ** p=≤0.01, *** p=≤0.001.

**Figure 3.**
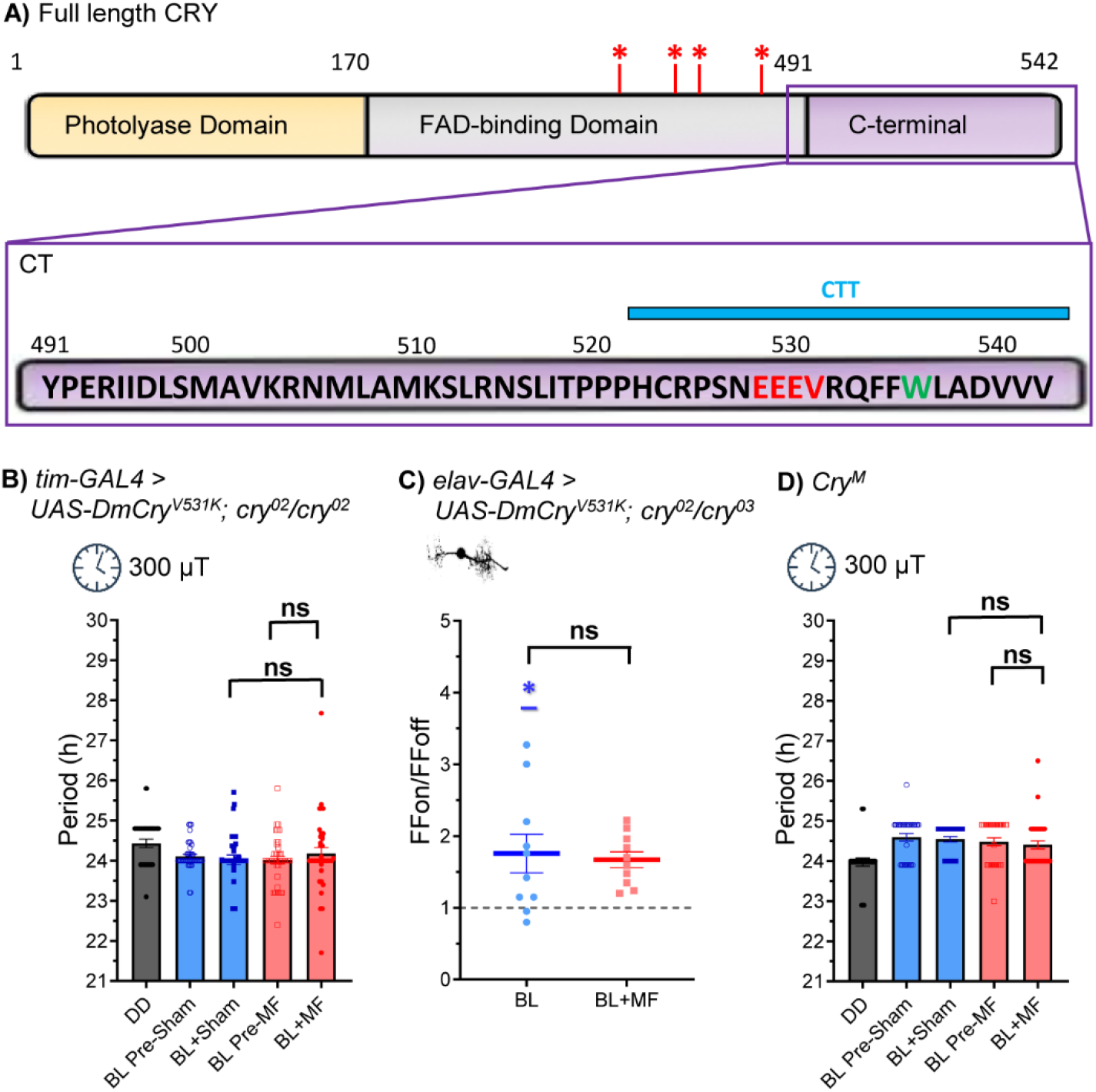
Integrity of the CTT is required for magnetosensitivity. (A). Schematic of the domain structure of CRY including the C-terminal (CT, aa 491-542) and CTT (aa 521-542). The four Trp residues, presumed essential for the canonical RPM, are indicated by red asterisks. A putative PDZ binding site (EEEV 528-531, shown in red) was mutated (V531) CRY^V531K^. The terminal Trp (W536) mutated in Luc-CT^W536F^ is shown in green. (B). CRY^V531K^ expressed in clock neurons *(tim-Gal4)* does not support magnetosensitivity in the circadian period-shortening assay. A 2-way ANOVA revealed no significant main, or interaction effects (interaction, F_1,52_=0.09, p=0.77, Extended Data Fig.6a). (C). Expression of CRY^V531K^ in aCC neurons is sufficient to support BL sensitivity (p=0.017, t_(9)_=2.934, n=10, Extended Data Fig.6b) but not a further potentiation in BL+MF (100 mT, p=0.768, t_(18)_=0.299, n=10). (D) CRY^M^, a truncated CRY mutant lacking the terminal 19 amino acids, including the PDZ binding motif (528-531), also fails to support a sensitivity to a 300 μT MF, (3Hz, F_1,122_=0.021, p=0.89, Extended Data Fig.6c). Blue asterisks represent significance values for aCC before *vs*. during BL exposure (paired t-test, same cells); ns (not significant) represent comparisons of BL *vs* BL+MF (unpaired t-test, different cells). ns p=>0.05, * p=≤0.05, ** p=≤0.01, *** p=≤0.001.

The Trp residue in CRY-CT has not been implicated in the canonical RPM. However, theoretically it is capable of generating a RP with free FAD reminiscent of the interaction between Flavin mononucleotide and the surface Trp of lysozyme^32^. Thus, we substituted this residue for a redox-inactive phenylalanine (i.e., W536F). Expression of Luc-CT^W536F^ was sufficient to lengthen circadian period in BL *vs.* DD (24.47 *vs*. 23.95 h, respectively, p=0.0017) indicating it supports circadian light-responsiveness. A 2-way ANOVA revealed a significant interaction between the pre/exposure and MF/sham treatment (F_1,198_=5.1 p=0.025, Fig.1Di). Expression of this variant also shortened the free-running circadian period when exposed to a MF (300 μT, 3 Hz) compared to their pre-exposure and to the sham exposed flies (p=0.023, p=0.015, respectively, Fisher LSD test, whilst the stringent Newman-Keuls test narrowly misses significance for both comparisons (p=0.063, p=0.074, Extended Data Fig.2c-e). Expression of Luc-CT^W536F^ also supported a strong (2-fold) BL response on AP firing in aCC and, again a significant, albeit more variable potentiation in BL+MF (Fig.1Dii, 2.69-fold, p=0.03, 2-way ANOVA replicates as a factor, Extended Data Fig.1c). That Luc-CT^W536F^ does not obliterate a MF response argues against a significant role for a hypothetical RP between W536 and FAD. Indeed, the weaker MF-response may be structural in origin^33^. An arginine (R532) in close proximity may form a cation-π interaction with W536 to stabilise an alpha helical conformation^34^ that would be disrupted by the W536F substitution, yet the MF effect is still detectable.

### Free FAD supports magnetoreception

The fact that Luc-CT^W536F^ is sufficient to support magnetosensitivity implies that a different, non-CRY, RP is involved. In this regard, it is notable that free FAD is capable of generating a magnetically sensitive RP *via* intramolecular electron transfer^7,35^. To explore this, we supplemented additional FAD to aCC *via* the internal patch saline. Increasing the concentration of FAD (10 to 50 μM, in the patch pipette) potentiates the efficacy of Luc-CT to mediate BL-dependent increases in AP firing (Fig.2A, R^2^=0.71, p=0.034), an effect that is enhanced in the presence of BL+MF (100 mT, p=0.015). Significantly, MF potentiation is by a fixed proportion relative to BL at each FAD concentration tested (evidenced by equal gradients of lines of best fit). This is a prediction of the RPM, providing biological saturation is not limiting, the proportional magnetically-induced change should remain constant^36^. In the absence of the Luc-CT construct, FAD (up to 50 μM) induced a weak, but significant, BL response (p=0.03, Fig.2B, Extended Data Fig.4a); however, no potentiating effect was observed in BL+MF.

FAD autoreduction occurs following electron transfer from the adenine side chain to the photoexcited isoalloxazine, generating an intramolecular RP^7,37^. To test whether such autoreduction is likely to support magnetosensitivity, we introduced riboflavin to the cell via the internal patch saline. Although riboflavin contains the same isoalloxazine chromophore and can populate photoexcited triplet states^38^, it lacks an adenine diphosphate side chain (Extended Data Fig.5) and is thus unable to generate the same intramolecular RP^39^. Riboflavin (50 μM), in the presence of Luc-CT, supported a BL effect (~1.94-fold, Fig.2C,), but there was no additional increase under BL+MF (100 mT, Fig.2C, p=0.9, Extended Data Fig.4b).

Our results are consistent with an interaction between FAD and Luc-CT, possibly in complex with other, unknown, molecules, which together may facilitate transduction of a magnetic field. Furthermore, our data suggest that molecules other than CRY are able to generate magnetically-sensitive RPs and produce a biological effect under appropriate conditions. *In vitro* spectroscopy has shown that BL photoexcited FAD generates RPs that are responsive to MFs^40^, and it appears likely that FAD is responsible for MF effects recently observed on cellular autofluorescence^35^. Thus, FAD (but not riboflavin) at higher concentrations may act as a magnetoreceptor. To test this, we recorded from aCC in a *cry* null background, which shows no overall BL or MF response (Extended Data Fig.3g). We observed that high levels of FAD in the internal patch saline (200 μM) were sufficient to support a BL-dependent increase in AP firing without need for Luc-CT (Fig.2D, 1.27-fold, Extended Data Fig.4c). Remarkably, this effect was potentiated in the presence of a MF (100 mT, Fig.2D, 1.84-fold, p=0.003). Cells supplemented with riboflavin (200 μM) showed an increase in AP firing in response to BL (Fig.2D, Extended Data Fig.4c) but did not show potentiation of the response in a MF (100 mT, p=0.67). That high levels of FAD alone are sufficient to support magnetosensitivity suggests that CRY-CT acts as an adaptor protein, possibly bringing photoactivated FAD close to its effector, HK. Proximity may allow HK to be activated directly by the resultant change in oxidative state that results from photoactivation of FAD. Very high levels of FAD negate this requirement. In the presence of CRY-CT the amount of photoactivated FAD required is presumably lower and, thus, more reflective of normal physiological amounts of this flavin.

### The integrity of the CTT is crucial for magnetic sensitivity

The less robust response from Luc-CT^W536F^ (Fig.1D) suggests that the integrity of the CTT might be important to its role in facilitating magnetosensitivity. The CTT of CRY also contains several linear motifs including putative PDZ-binding sequences^41^ *(e.g.,* EEEV 528-531, Fig.3A). PDZ proteins function as modular scaffolds that direct the cellular localisation of signalling molecules such as ion channels (for instance, Shaker K^+^) ^42,43^ and assembly of signalling partners (including CRY) into a ‘signalplex’ of the phototransduction cascade in the *Drosophila* eye^21,22^. To explore the importance of CTT structure and, specifically, to determine whether the putative PDZ-binding motif at residues 528-531 regulates magnetosensitivity, we mutated valine to lysine in position 531 (V531K)^41^ in full-length DmCry. Pan-circadian expression (*i.e.,* using the *tim-GAL4* driver) of DmCry^V531K^ in a *cry^02^* null background retained a circadian light sensitivity with a slight period shortening (Fig.3B, DD=24.51 *vs.* BL=24.20 h, p=0.005, Grubbs outlier test excluded a single very weakly rhythmic short period (20.3 h) fly in DD, see Methods), but failed to support a whole-organism behavioural response to MF (Fig.3B, interaction F_1,158_=0.55, p=0.33, Extended Data Fig.6a). Expression of DmCry^V531K^ in aCC showed the expected effect of BL on AP firing (Fig.3C, 1.76-fold, Extended Data Fig.6b). As in the circadian assay, this variant was unable to support magnetosensitivity in the aCC neuron (100 mT, Fig.3C, p=0.77). The loss of a MF effect, but retention of a BL response for DmCry^V531K^ is reminiscent of the CRYΔ mutant^3,4^, which lacks the CTT entirely. To further validate this result, we used the CRY^M^ mutation that lacks the final 19 AAs of the CTT^44^, including the putative PDZ domain centred around 531. This CRY mutation retained circadian light sensitivity but did not show a MF-induced period shortening as tested at both 300 μT, (3Hz, Fig.3D, F_1,122_=0.021, p=0.89, Extended Data Fig.6c), and 50 μT (3Hz, F_1,180_=0.3, p=0.6 Extended Data Fig.6d-e). These results confirm the CTT as a probable mediator of the MF response, where it likely serves to facilitate formation of protein complexes that transduce the magnetic signal.

### CRY4 of the migratory European robin mediates MF effects in *Drosophila*

Finally, recent *in vitro* spectroscopic studies have suggested that CRY4, encoded in the genome of the European robin, *E. rubecula* a migratory songbird^45^ may represent the magnetoreceptor responsible for long-distance navigation in this species. We generated a *UAS-ErCry4* transgene and expressed it in the clock neurons of the fly. We observed significant period-shortening on exposure to a MF compared to sham at 300 μT MF (3Hz, 2-way ANOVA F_1,238_=4.4, p=0.036, Fig.4A), or 50 μT (3Hz, 2-way ANOVA, interaction F_1,237_=4, p=0.047, Fig.4B, Extended Data Fig.7a-f). Expression of ErCry4 in aCC was also sufficient to render the cell BL sensitive (1.8-fold, Fig.4C, Extended Data, Fig.7g) and sensitive to an external MF (100mT, 2.94-fold, p=0.046).

**Figure 4.**
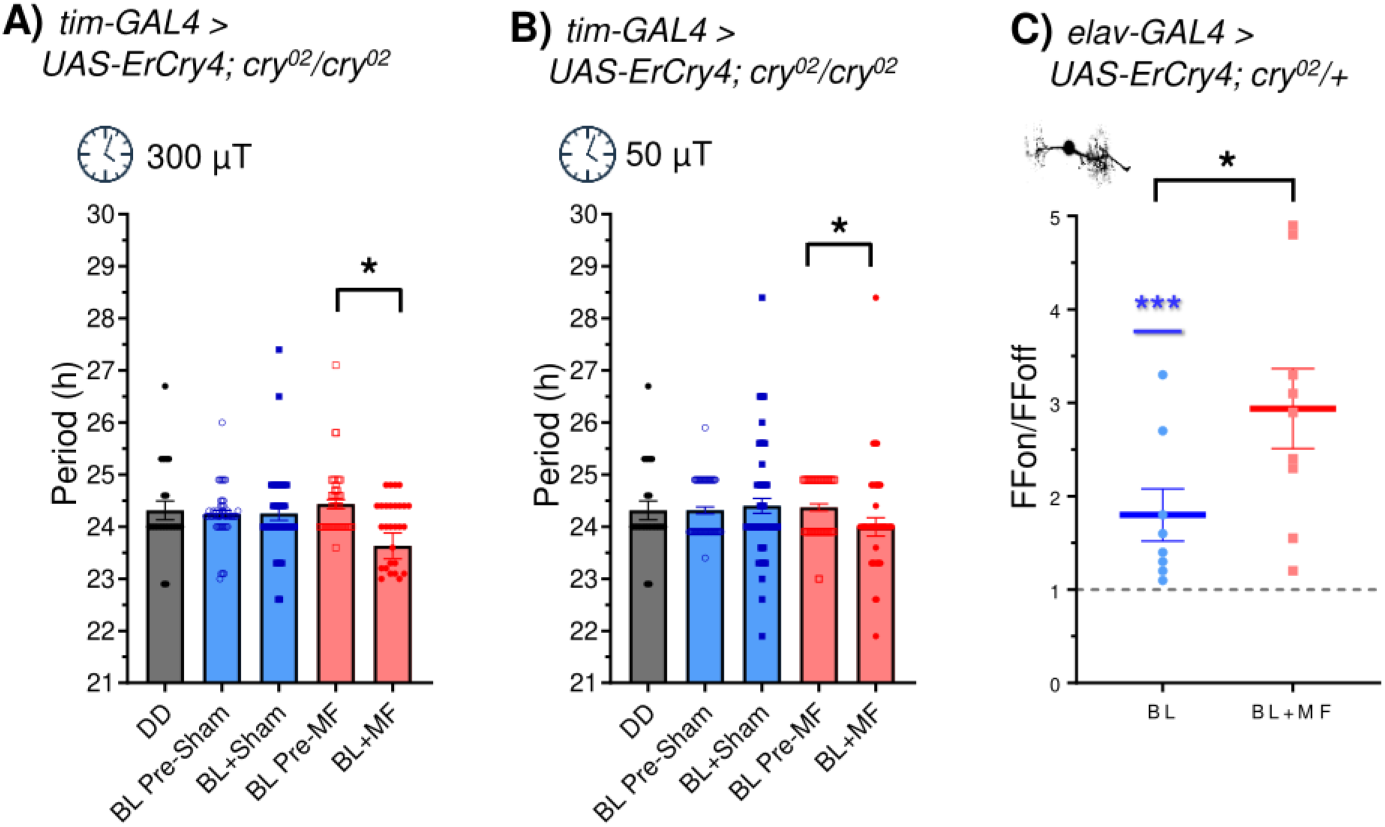
ErCry4 is sufficient to support MF-sensitivity in *Drosophila*. (A). Expression of European Robin (*Er)Cry4* in *Drosophila* clock neurons (tim-GAL4) results in a significant period shortening in the presence of a 300 μT MF (3Hz, 2-way ANOVA interaction F_1,238_=4.4, p=0.036). (B). Period shortening is also present in a 50 μT MF (3Hz, F_1,237_=3.98 p=0.047). (C) Relative firing frequency recordings of the aCC motoneuron, expressing ErCRY4, in BL vs. BL+MF. BL exposure alone leads to an increase in the number of APs (1.8-fold, (t_(7)_=3.6, p=0.0088, n=8, Extended Data Fig.7g), and the co-presence of a MF (100 mT) induces a significantly greater firing-fold change of 2.94 (BL vs BL+MF, t_(15)_=2.17, p=0.046, n=9). MF-potentiation between the two groups was tested by unpaired t-tests.

### Conclusions

We have observed that contrary to several reports^2,14^, but not others^3^, full length CRY may be sufficient, but is not strictly necessary, to mediate magnetosensitivity. The expression of the C-terminal 52 residues of CRY is sufficient to support magnetosensitivity in both single neuron and whole animal assays. Our results challenge the canonical CRY-dependent RPM model of animal magnetoreception (based on the requirement for full-length CRY, including FAD binding and the Trp chain), but are nevertheless consistent with a RPM. It remains unknown if Luc-CT binds FAD directly. Yet, the Luc-CT response is potentiated by increasing the cytosolic availability of FAD, a common biological redox cofactor, implying that redox reactions are at the core of magnetosensitivity^46^. We cannot exclude that alternative RPs that are not directly photochemically-generated, may also contribute to magnetoreception. This would be consistent with a growing list of examples reporting RP mediated magnetoreception in darkness^30,47–49^. The synergistic interaction between Luc-CT and free FAD argues the former facilitates formation of a complex that enables the transduction of a magnetically-derived signal by the latter. Moreover, free FAD itself can mediate a magnetic response *in vivo* but at high, non-physiological, levels. We interpret these results to suggest that evolution has shaped the defining element of CRY, its CT, to bring the RP to the proximity of cellular effectors (for instance, HK). Thus, through protein-protein interactions, CRY can ‘potentiate’ the weak activity of the geoMF on any associated RP. In this regard, the primary role of CRY would be that of a magnetotransductor and then of a magnetoreceptor.

The unexpected observation that robin ErCry4 can also mediate MF effects in *Drosophila,* in both circadian and electrophysiological assays, argues that the fly is an excellent tractable model system to dissect the molecular component of magnetoreception. Why might flies have a magnetic sense, given that they do not navigate/migrate in the same way as birds? While our circadian phenotype is somewhat contrived and seeks to use a sensitised CRY background (dim constant BL), which would provide the best opportunity for observing any MF effects, dCRY is not only a circadian photoreceptor; it also mediates geotaxis behaviour^50^. Independent studies have revealed that geotaxis shows a dCRY-dependent magnetosensitivity^51,52^. Remarkable results have suggested that flies exposed to a MF as embryos are ‘imprinted’ on the MF in which they develop and as adults they prefer to forage with downward movement in their home MF^53^. As *D. melanogaster* is well-known to forage/mate/oviposit on rotten fruits that are usually found at ground level, this geotactic magnetic sense would appear to have fitness value.

In conclusion, our observations suggest an ancient and ubiquitous effect of MFs on biological RPs. Through CRY, evolution has optimised such an effect by bringing together two functions, ‘receptor’ and ‘transductor’ that are required for magnetosensing but not necessarily as parts of the same molecule. That *Drosophila* (and other non-migrating animals) can sense external magnetic fields has been reported by many independent, groups^54^. This is seemingly reflective of the physiochemical properties of flavins such as FAD to form RPs. In animals that do navigate, this mechanism has presumably been adapted to underpin this behaviour. The underlying physiochemical properties of CRY-dependent magnetosensitivity appear to be shared across navigating and non-navigating animals.

## Materials and Methods

### Fly stocks

For larval aCC recordings, embryos were raised at 25°C in a 12:12 light / dark cycle until 3^rd^ instar wall climbing larvae (L3) emerged, these were then kept in darkness through the day of recording to minimise light dependent CRY degradation. Recordings were conducted between circadian time hours: 2-10. Flies were maintained on standard corn meal medium at 25°C. The driver line *elavC^155^-GAL4;*; *cry^03^* was obtained from Bloomington Stock centre (#BL458) and crossed into a *cry^0^* background as described^55^. *cry^0^* flies were obtained from Bloomington Stock centre (#BL86267), *tim-GAL4; cry^02^* and *UAS-cry; cry^02^* are already described^3,56^. *cry^M^* (kindly supplied by Dr David Dolezel (Institute of Entomology, Czech Academy of Science) has a stop codon inserted at AA523 and lacks the final 19 AAs of the CTT, which includes the putative PDZ binding motif at 531 (see Fig.3A). The generation of novel transgenic flies for this study are described below.

### Molecular Cloning of Luc-CT

Luciferase CDS was cloned from the *UAS-Luc-CRY* fly line^18^ and subsequently amplified with the following primers to include overhangs compatible for the NEB Gibson Assembly assay

F: *TATCCTTTACTTCAGGCGGCCGCATGGAAGACGCCAAAAACATAAAGAAAGG;*

R: *TCCGGATACTCGAGCACGGCGATCTTTCCGCCC.*

The CT portion of *cry* was produced by gene synthesis (GeneArt, ThermoScientific) based on the original *GFP-CT* construct^3^. *CT* was designed to include 5’ and 3’ overhangs compatible with subsequent NEB Gibson Assembly assay

(Seq:*AAAGATCGCCGTGCTCGAGTATCCGGAGCGGATCATTGATTTGTCCATGGCCGTGAAGCGCAACATGCTGGCCATGAAGTCCCTGCGCAACAGCCTGATCACCCCCCCACCACATTGCCGCCCCAGCAATGAGGAGGAAGTGCGCCAGTTCTTCTGGCTGGCCGATGTGGTGGTGTAATCTAGAGGATCTTTGTGAAGGA*).

pJFRC2-10XUAS-IVS-mCD8::GFP (Addgene #26214) plasmid was digested with *NotI* and *XbaI.* A Gibson Assembly assay was performed to ligate pJFRC, *Luc* and *CT* in a single step reaction. A Myc tag was produced by annealing single Oligos designed using The Oligator software (14avidson.edu) and ligated 5’ of Luc-CT. Briefly, 5 μM of Oligos

#### 47-mer Top1

*5’-GATCTCACAATGGAACAGAAGCTGATCTCCGAGGAGGACCTGGGCGC*

#### 47-mer Bottom1

*5’ GGCCGCGCCCAGGTCCTCCTCGGAGATCAGCTTCTGTTCCATTGTGA*

Resulting in 5’*BglII* and 3’*Notl* over-hangs once annealed, were diluted in 1X annealing buffer (0.1 M NaCl; 10 mM Tris-HCl, pH 7.4), boiled in 500 μl of H2O for 10 minutes and left overnight to cool down to room temperature to hybridise. The pJFRC2-10XUAS-Luc-CT plasmid was then digested with *BgIII* and *NotI*. The fragment encoding the Myc tag was then ligated using standard methods. After sequence validation, the plasmid was injected into the *y w, M(eGFP, vas-int, dmRFP)ZH-2A; P{CaryP}attP40* (Stock 13-40, Cambridge University Fly facility) using the Phi31 integrase system for insertion. The resulting transformants were subsequently backcrossed into the *w^1118^* background for 7 generations. The Luc-CT^W536F^ transgene was generated by gene synthesis (Eurofins, Ebersberg, Germany) and sub-cloned into pJFRC-MUH via 5’*NotI* and 3’*XbaI* restriction sites. Transgenic injections for Luc-CT^W536F^ were carried out by Manchester University Fly Facility using the same *y w M(eGFP, vas-int, dmRFP)ZH-2A; P{CaryP}attP40* line (Stock 13-40, University of Cambridge Fly Facility). The *ErCry4* transgenic was generated by gene synthesis (NBS Biological Ltd., Huntingdon, UK) and sub-cloned into pJFRC-MUH via *NotI* and *KpnI* restriction sites. Injection was carried out by Bestgene, California, USA) into *y w; PBac{y[+]-attP-3B}VK00002* (Bloomington no. 9723) via the Phi31 integrase system.

*HA-cry^V531K^* (containing a HA tag at the N-terminal of the encoded protein) was already available as a clone in the yeast plasmid pEG202^41^. It was released by *EcoRI-XhoI* digestion and sub-cloned into pUAST^57^. Transgenics were produced by P-element transformation by the University of Cambridge Fly Facility using the line S(6)1 inserted on chromosome 2.

### Electrophysiology

The experimenter was blinded to genotype during both recordings and subsequent data analysis. L3 larvae were dissected under extracellular saline as described^58^ with the only modification being a red filter applied to both the dissecting light and compound microscope to minimise CRY degradation prior to, and during, recordings. Thick-walled borosilicate glass electrodes (GC100F-10; Harvard Apparatus, UK) were fire polished to resistances of 10 - 15 MΩ. Recordings were made using a Multiclamp 700B amplifier controlled by pCLAMP (version 10.4) *and* Digidata 1440A analog-to-digital converter (Molecular Devices, CA, USA). Only cells with input resistance of ≥500 MΩ were used. Traces were filtered at 10 kHz and sampled at 20 kHz. The extracellular saline solution contained the following (in mM): 135 NaCl, 5 KCl, 4 MgCl_2_.6H_2_O, 2 CaCl_2_.2H_2_O, 5 TES, and 36 sucrose, pH 7.15. The intracellular patch solution contained the following (in mM): 140 K-D-gluconate, 2 MgCl_2_.6H_2_O, 2 EGTA, 5 KCl, and 20 HEPES, pH 7.4. KCl and CaCl_2_ were from Fisher Scientific (UK); sucrose was from BDH (UK); all remaining chemicals were from Sigma-Aldrich (UK). Mecamylamine (1 mM) was applied to all preparations to isolate the aCC motoneurons from excitatory cholinergic synaptic input. For recordings supplemented with additional FAD or riboflavin (Sigma-Aldrich, UK), dilutions were made up in intracellular saline and kept in the dark.

### Photoactivation and Magnetic Field application

Light stimulation was supplied by a blue LED (470 nm, Cairn Research, UK) at a power of ~2.2 mW/cm^2^, a value used previously to stimulate CRY^59^. Each cell was injected with a variable amount of constant current until threshold potential was reached and the neuron was allowed to settle, for some minutes, until AP firing was stable at ~5-7 Hz. Once a stable firing rate was achieved, each neuron was recorded for at least 20 s before exposure to BL illumination for 30 s. No change to AP firing rate was observed without BL illumination. Magnetic exposure was provided by two NdFeB static magnets mounted around the preparation at a distance that provided a MF of 100 (± 5) mT. Field strength was measured using a 5180 Gauss/Tesla Meter (F.W. Bell. USA). This method is essentially identical to that used previously^4^.

### Statistical analysis of electrophysiological recordings

A D’Agostino & Pearson analysis showed our data to be normally distributed, thus parametric tests were applied in all cases. Data are shown as mean ± SEM. To determine BL sensitivity a paired t-test (two-tailed) was used to compare the number of APs a neuron fired in the 15 s following light stimulation *vs.* the number of APs in the preceding 15 s before light exposure. For comparison between BL and BL+MF the number of APs in the 15 s preceding and proceeding BL or BL+MF exposure was used to determine the firing-fold change (FF_on_/FF_off_) for each cell. Statistical significance for MF potentiation against BL effect alone was determined using an unpaired t-test (two-tailed) to compare the firing-fold change for the BL dataset *vs.* BL+MF dataset. Where multiple genotypes/conditions were tested simultaneously, a 2-way ANOVA with Newman-Keuls *post-hoc* testing was used. For the FAD dose response curve, a BL effect of [FAD] was determined by linear regression fitting and significance determined using an ANCOVA model. Average MF potentiation of the Luc-CT FAD dose response (Fig.2A) was determined based on the intercept of the Y axis.

Unpaired t-tests (two-tailed) were also applied in the Extended Data figures to compare the number of APs in the ‘before’ BL±MF conditions, as well as to BL and BL+MF exposures. Control lines were also compared to their respective experimental genotype by both 1-way ANOVA (with BL and BL+MF recordings separated) and by 2-way ANOVA. Raw data are reported in the Extended Data.

### Behavioural analyses and statistics

Circadian locomotor activity was recorded using a *Drosophila* Trikinetics Monitor 2 (Waltham, MA, USA)^3^. To test the effects of MF on the free-running circadian period of locomotor activity, we used a modified version of the Schuderer apparatus, which consists of two independent double wrapped coils placed inside two μ–metal boxes within a commercial incubator. The shielded, four quadratic Helmholtz coil systems produce a homogenous, linearly polarized *B* field with perpendicular orientation to the horizontal plane of the Trikinetics monitors. Each coil is formed with a pair of wires, with the current passing in the same direction through both wires for MF exposure but in opposite directions to provide a sham exposure condition (0 T). A computer randomly assigns the MF and Sham exposed chambers and the experiment is performed blinded^17^.

One to three day old flies were first entrained at 20 °C in the apparatus under a dim BL: darkness 12 h cycle (BL: DD = 12:12) for three full days, before being pre-exposed to continuous BL for 7 days, followed by exposure to BL+MF or BL+Sham for a further 7 days^3^. Thus, there were 4 measurements, the pre-exposure (BL) period of flies that were subjected to a MF or sham, plus the exposure period for both (BL+MF and BL+Sham). A fifth control condition examined the period of Luc-CT*; cry^02^* in DD without exposure. All experiments were performed using a low frequency 3 Hz field at 300 μT and dim BL at 0.15 - 0.25 μW/cm^2^, wavelength 450 nm, 40 nm broad range (RS Components, UK). The driver *tim-*GAL4 was used to express UAS-CRY transgenes as previously described^3^.

Rhythmicity and period were determined using spectral analysis employing a MatLab-based version of the BeFly program^60^. Statistical analysis of period was performed using ANOVA with either Statistica (Statsoft, CA, USA) for factorial analyses or Prism (Graphpad) for 1-way ANOVA. Although there was a clear prediction that Luc-CT flies would have a shorter period under a MF^3^, we nevertheless used the stringent Newman-Keuls *post-hoc* test to compare groups after factorial ANOVA buttressed by the more liberal Fisher LSD test for the circadian Luc-CT^W(536)F^ results. To compare the DD periods with those from the BL pre-exposure conditions we used an unpaired t-test. Circadian data were first tested using a Grubbs outlier test (GraphPad Prism, alpha =0.01 two-sided, Z=5.3). One datum from the DD data of CRY^V(531)K^ which represented the least robust single period in the dataset with an anomalous period of 20.3 h (8 sd away from the mean) was identified and removed.

## Acknowledgements

We thank Matt Hemsley and John Hares for the cloning and production of CRY-CT mutants and flies. ARJ thanks Ines Camacho and Mike Shaw for useful feedback on the manuscript. This work was supported by funding from the Leverhulme Trust to RAB, and ARJ (RPG-2017-113) and to RAB from BBSRC (BB/V005987/1) and to CPK/ER from BBSRC (BB/V006304/1). Work on this project benefited from the Manchester Fly Facility, established through funds from the University and the Wellcome Trust (087742/Z/08/Z). ARJ thanks the National Measurement System of the Department for Business, Energy and Industrial Strategy for funding. GF, CNH, ER and CPK acknowledge funding from the Electromagnetic Field Biological Research Trust.

