## Extended Data files for "Essential elements of radical pair magnetosensitivity in *Drosophila*"

Extended Data Figure 1

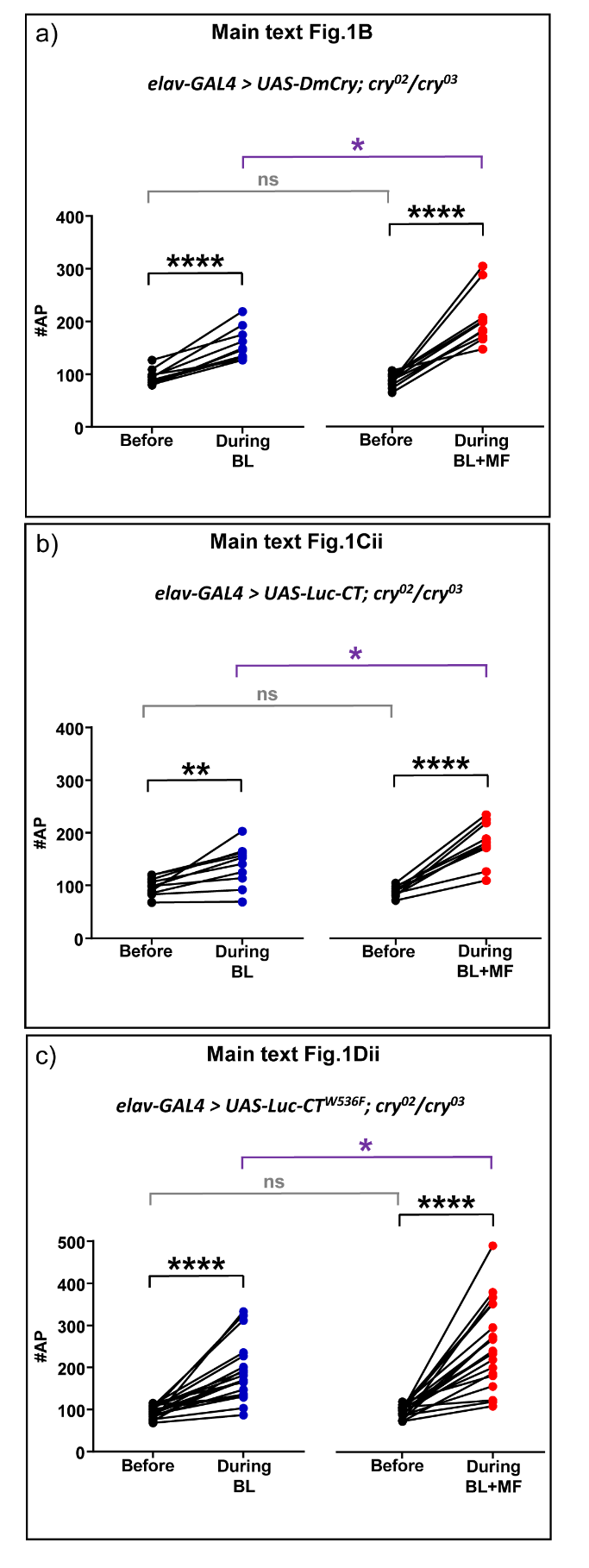

**Extended Data Figure 1.** Supporting electrophysiological data for main text Fig 1.

Raw action potential (AP) counts for neurons expressing (a) DmCry, (b) Luc-CT or (c) Luc-CT^W536F^, recorded in the 15 s before and 15 s during exposure to BL or BL±MF. This data was used to derive firing-fold change reported in main text Fig.1. Paired t-tests were used to compare AP counts before *vs*. during for cells exposed to BL (left hand graph) or to BL±MF exposure (right hand graph). MF-potentiation between the two groups was tested by unpaired t-test (different cells). The figure number at the top of each panel represents the main text figure supported. ns p=>0.05, * p=≤0.05, ** p=≤0.01, *** p=≤0.001. See main text Fig 1 for FF_on_/FF_off_ ratio comparisons.

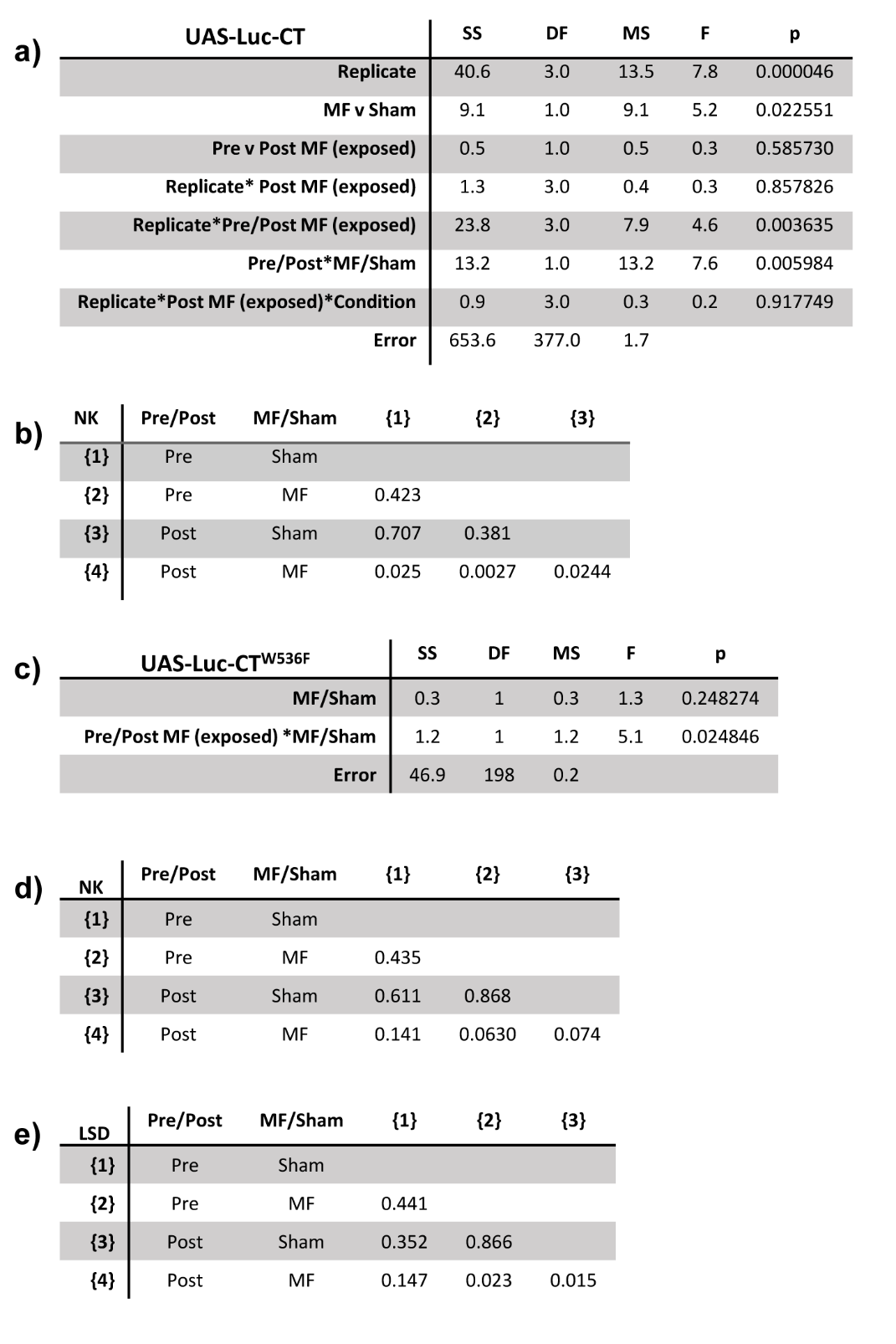
Extended Data Figure 2

Extend Data Figure 2 continued:

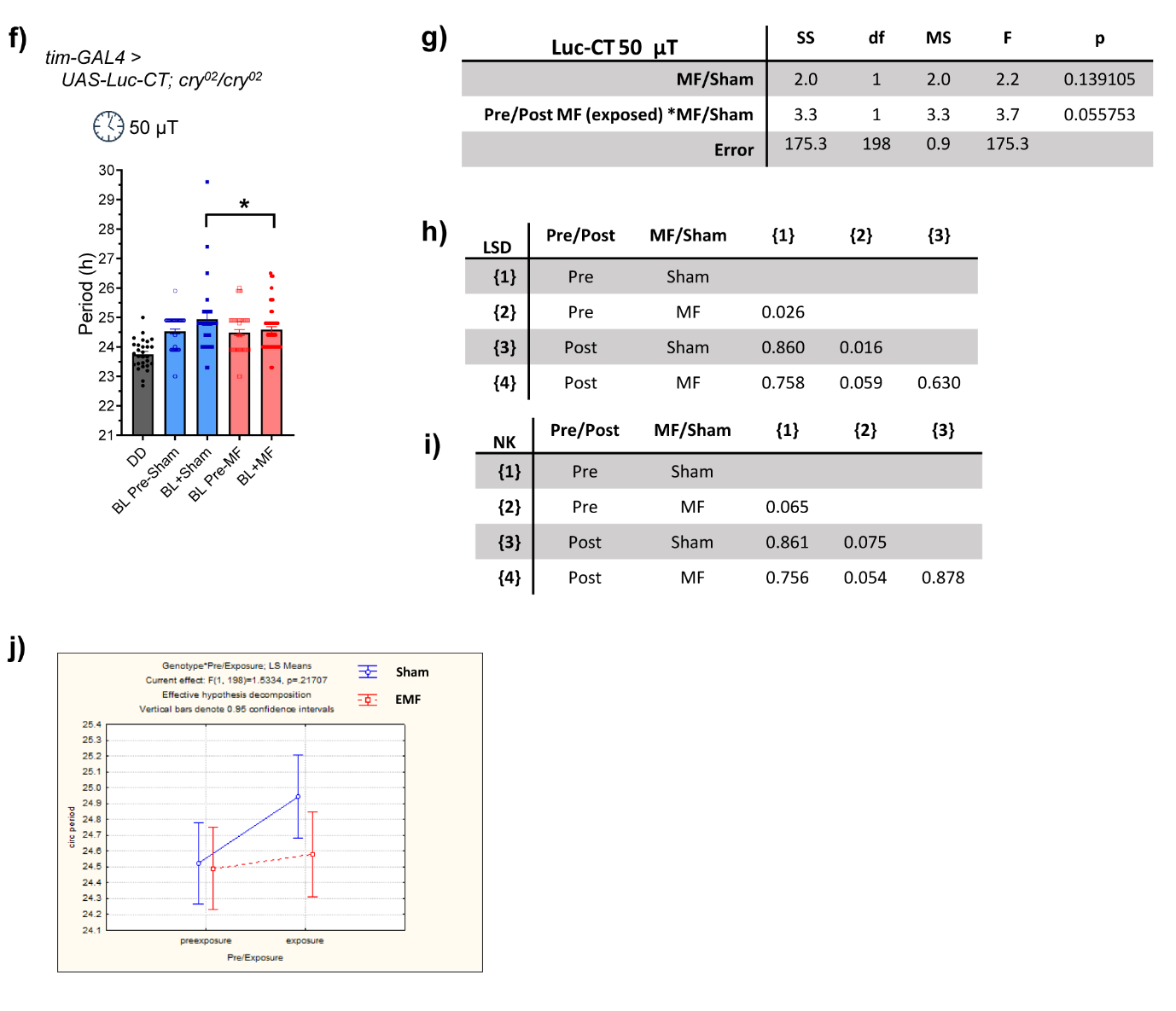

**Extended Data Figure 2.** Supporting circadian data for main text Fig 1.

(a). Table of 3-way ANOVA of BL conditions reveals a significant Sham / MF x pre / post-exposure interaction (F_1,377_=7.6, p=0.006) with MF (300 µT) exposed flies showing a significantly shorter period than sham exposed flies. (b). Table of Newman-Keuls *post-hoc* comparisons from a. (c). Table of a 2-way ANOVA of Luc-CT^W536F^ circadian experiment reveals significant interaction (F_1,198_=5.1, p=0.025) (d) Tables of Newman-Keuls *post-hoc* and (e) Fisher LSD comparisons reveal a MF field effect on Luc-CT^W536F^ circadian period in the latter test. The relevant comparisons in the more stringent NK test approach significance. (f). Exposure of Luc-CT to a 50 µT (3Hz) MF results in a small and marginal reduction in period compared to sham (g). 2-way ANOVA table (interaction F_1_,_198_=3.7, p=0.056) on circadian period for Luc-CT at 50 µT. (h). LSD and (i). Newman-Keuls *post-hoc* tests reveal a marginal effect (p=0.059, 0.054 respectively) on period shortening. (j). Box and whisker diagram (95% confidence limits) showing a relative period shortening under MF (EMF-red) compared to the BL+Sham (blue) exposure.

Extended Data Figure 3
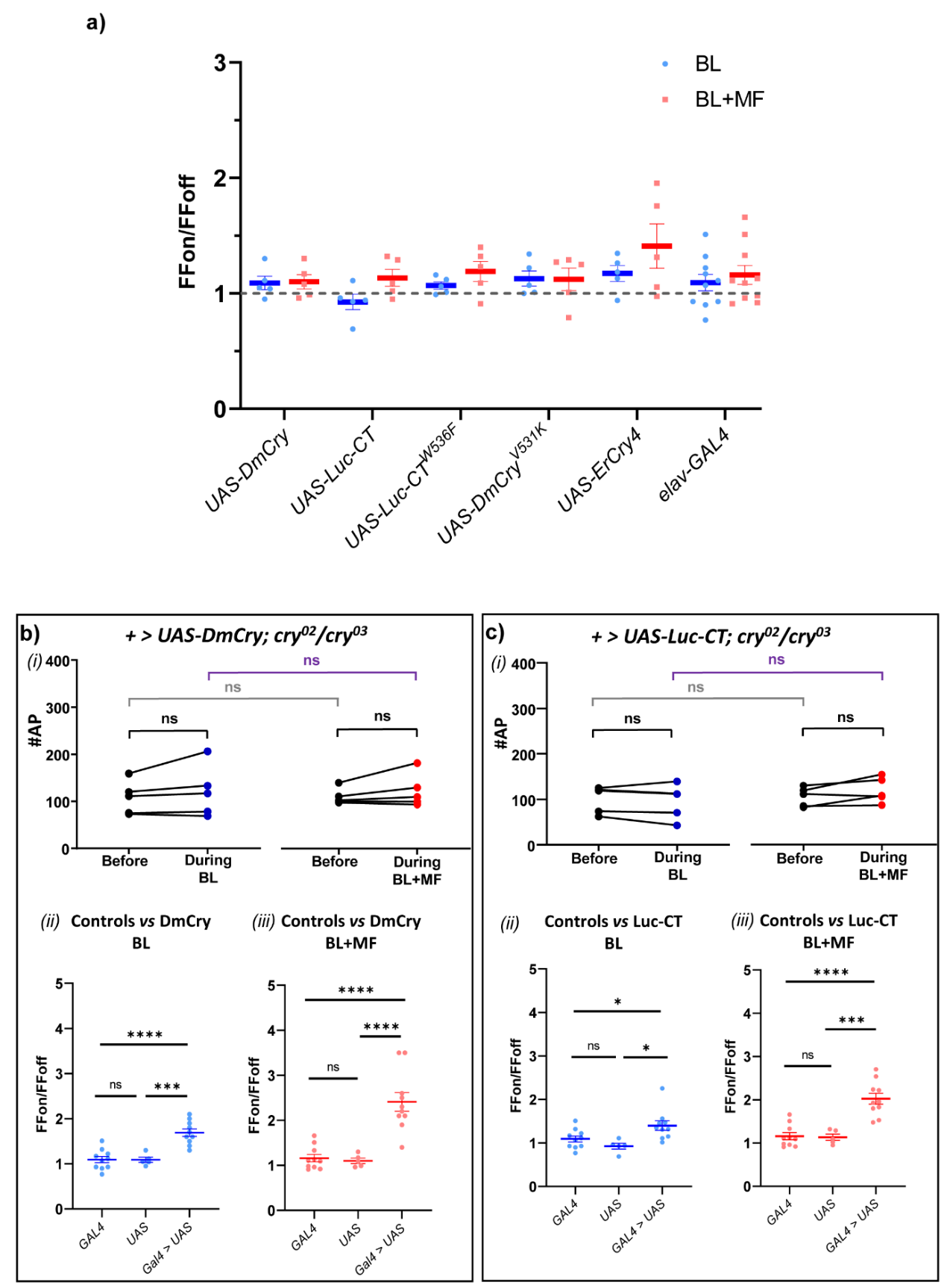

Extended Data Figure 3 continued:

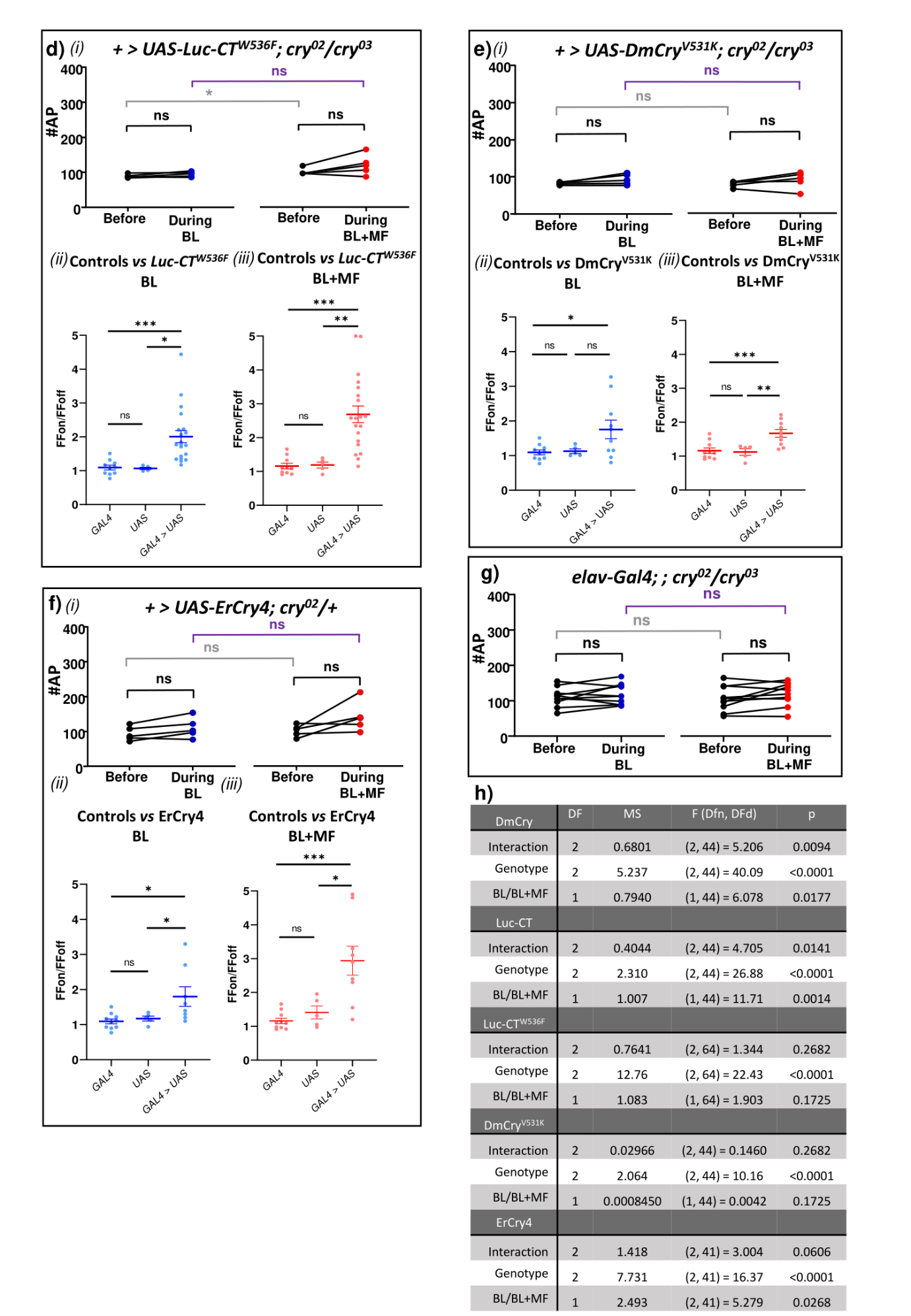

**Extended Data Figure 3.** Supporting electrophysiological data for main text Fig 3.

(a). Averaged data for parental (control) *Gal4* and *UAS* genotypes separately in a CRY null mutant background. Without the *Gal4* and *UAS* elements combined the CRY transgene (under *UAS* control) is not expressed and no significant BL or BL+MF response is seen. A 2-way ANOVA of the controls in both BL or BL+MF showed no significant interaction (F_(4,50)_=0.52, p=0.719). (b-g). Raw AP counts *(i)* for each aCC neuron recorded in the 15 s before *vs*. the 15 s during BL or BL±MF exposure for expression of the respective CRY transgene (paired t-tests, two-tailed). As an additional comparison of MF effect, unpaired t-tests (two-tailed) were used to determine significant differences between raw AP counts for both ‘before’ exposure conditions (grey line), and for during BL *vs*. BL+MF exposure (purple line). These comparisons revealed no significant differences in AP counts between the ‘before’ conditions for BL and BL+MF, apart from (di) *+ >UAS-Luc-CT^W536F^; cry^02^/cry^03^*. The co-presence of a MF (100 mT) had no effect on the number of APs in BL+MF compared to those exposed to BL alone. A one-way ANOVA is also shown for each control, comparing the expressed transgene with control parental UAS-transgene (*cry^02^/cry^03^*) or *elav-Gal4 driver; ; cry^02^/cry^03^* for both BL *(ii)* and BL+MF *(iii)*. These reveal no significant differences between the parental UAS lines and *elav-Gal4; ; cry^02^/cry^03^*, and a significant increase in firing-fold change for all expressed CRY transgenes compared to *elav-Gal4; ; cry^02^/cry^03^* or their respective UAS control, with the exception of (e*iii*) *elav-Gal4 >DmCry^V531K^; cry^02^/cry^03 ­­^*compared to *+ >UAS-DmCry^V531K^; cry^02^/cry^03^* in BL. (h). A 2-way ANOVA comparing each UAS-transgene, and the *elav-Gal4; ; cry^02^/cry^03^* driver line compared to the expressed CRY transgene in BL and BL+MF was also performed for each CRY variant. Although the 2-way ANOVA for Luc-CT^W536F^ reports no significant interaction, there is a significant genotype effect and Newman-Keuls *post-hoc* tests revealed a significant effect of MF compared to BL alone (main text Fig.1Dii). *elav-Gal-4* driver n=10, UAS-transgenic n=5 for both BL and BL+MF. ns p=>0.05, * p=≤0.05, ** p=≤0.01, *** p=≤0.001.

Extended Data Figure 4

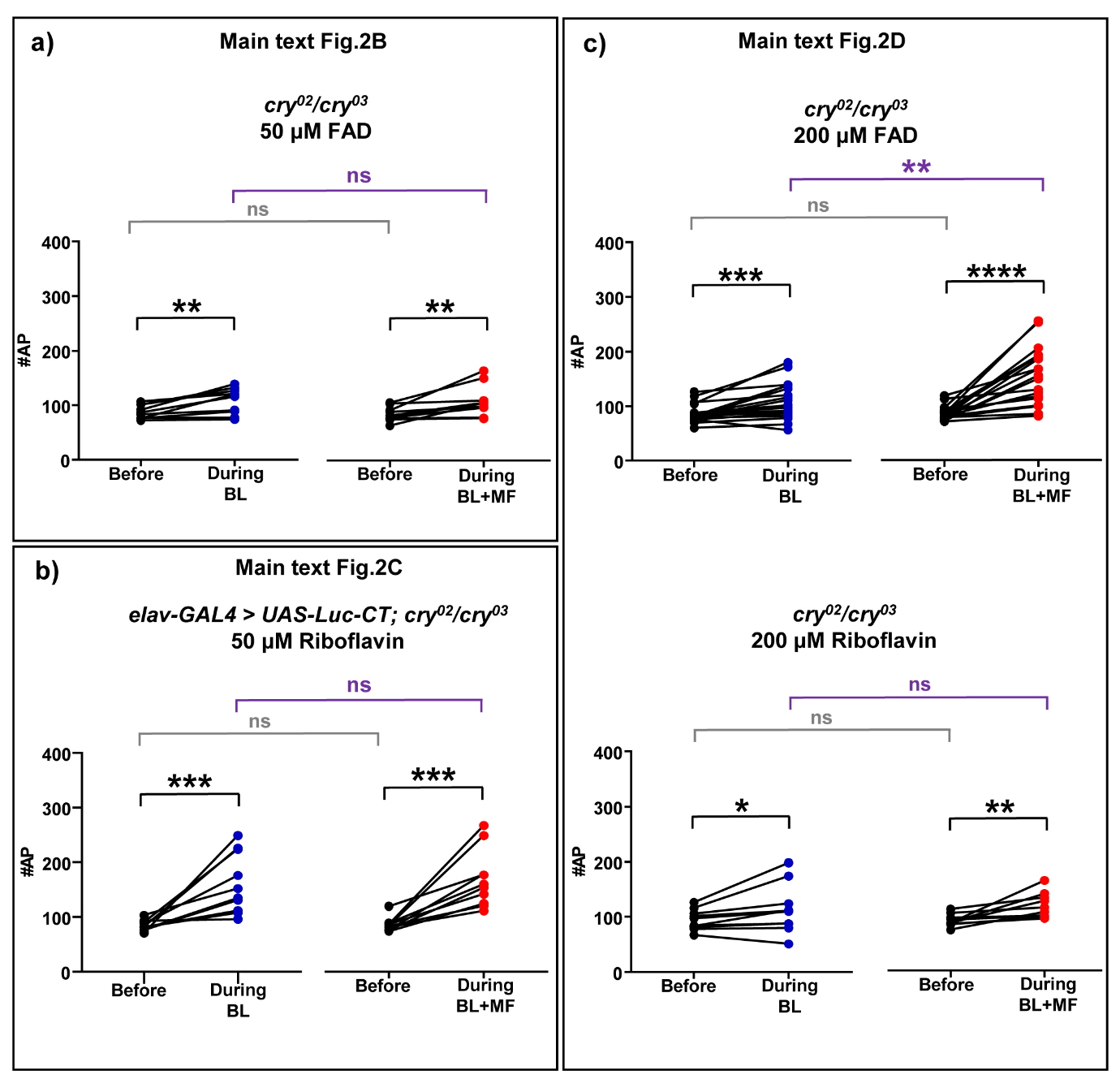
**Extended Data Figure 4.** Supporting electrophysiological data for main text Fig 2.

Raw AP counts for averaged data shown for flavin supplementation in main text Fig2.B-D. (a). *cry^02^/cry^03^* null cells supplemented with 50 µM FAD show a BL response, but no MF potentiation compared to BL alone (see main text). (b). Cells expressing Luc-CT supplemented with riboflavin (50 µM) show no MF effect. (c). Supplemented FAD and riboflavin (200 µM) to *cry^02^/cry^03^* null cells: FAD supports BL and BL+MF sensitivity, whilst riboflavin only supports BL sensitivity (see main text for BL and BL+MF comparisons). Paired t-tests were used to compare before *vs*. during for cells exposed to BL (left hand graph) or to BL±MF exposure (right hand graph). MF-potentiation between the two groups was tested by unpaired t-tests (different cells). All experiments have an n=10 for both BL and BL+MF, apart from the 200 µM supplementation to the *cry^02^/cry^03^* null (BL n=20, BL+MF n=19). ns p=>0.05, * p=≤0.05, ** p=≤0.01, *** p=≤0.001.

Extended Data Figure 5

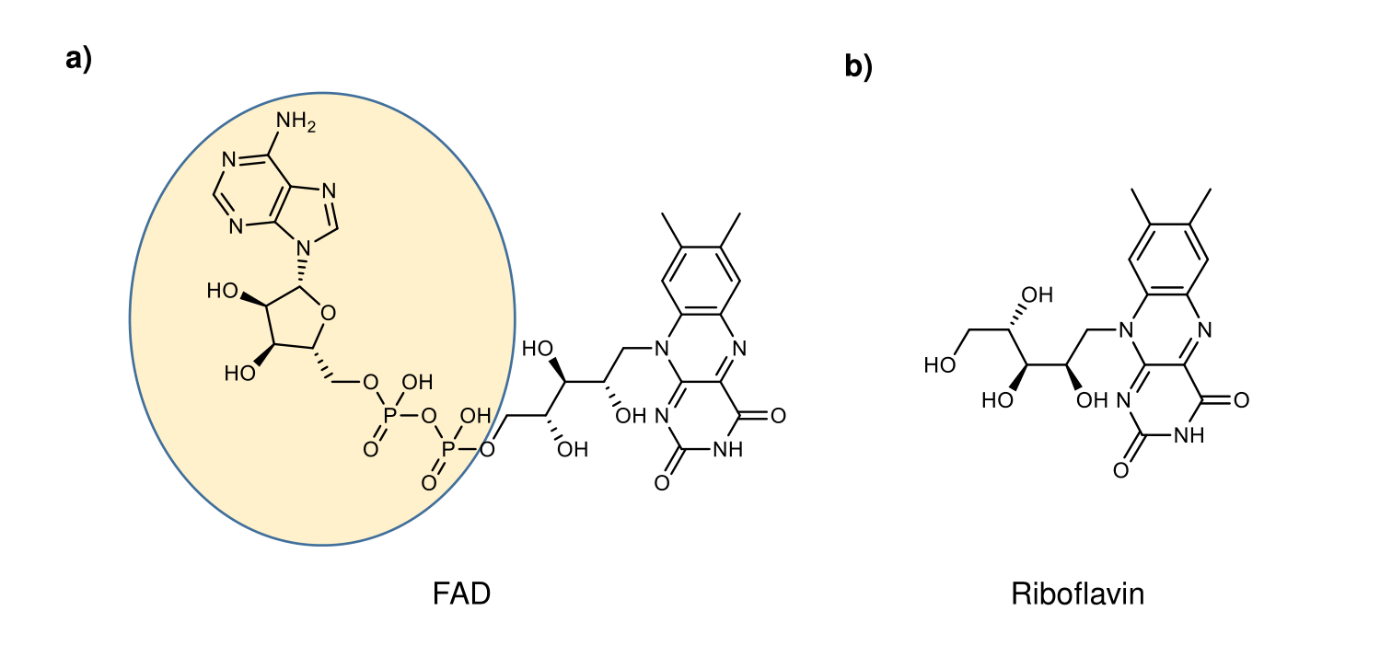

**Extended Data Figure 5.** Structures of FAD and Riboflavin.

The molecular structure of the two flavin chromophores. (a). Flavin Adenine Dinucleotide (FAD). Note the adenine diphosphate side chain of FAD (yellow oval), which facilitates the generation of an intramolecular magnetically sensitive RP. (b). Riboflavin, which is a metabolic precursor to FAD lacks the diphosphate side chain.

Extended Data Figure 6

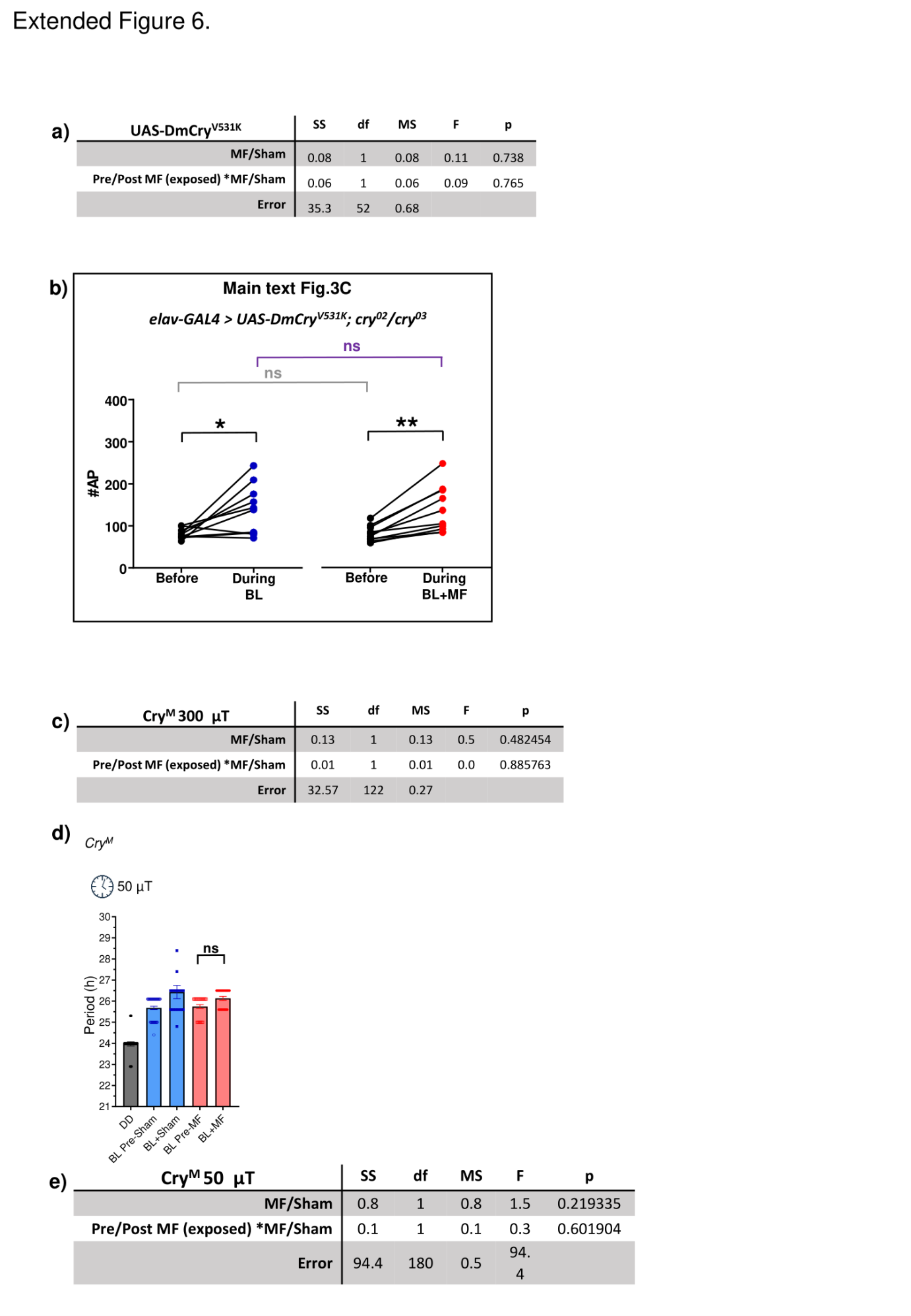

**Extended Data Figure 6.** Supporting circadian and electrophysiology data for main text Fig 3.

(a). Table of 2-way ANOVA of DmCry^V531K^ circadian period shortening shows no significant main nor interaction effects (F_1,158_=0.55, p=0.33). (b). Raw AP counts for each aCC neuron recorded, expressing DmCry^V531K^ in the 15 s before *vs*. the following 15 s during BL±MF, from which the average firing-fold change was derived for data reported in main text Fig.3C. Paired t-tests were used to compare before *vs*. during for cells exposed to BL (left hand graph) or to BL±MF exposure (right hand graph). MF-potentiation between the two groups was tested by unpaired t-tests (different cells). ns p=>0.05, * p = ≤0.05, ** p=≤0.01, *** p=≤0.001. (c). Summary of the 2-way ANOVA of CRY^M^, a protein encoded by the *cry^M^* allele of the endogenous *cry* locus that lacks the PDZ binding motif and fails to support period shortening in a MF (300 µT, 3Hz, interaction F_1,122_=0.021, p=0.89. (d). *cry^M^* flies do not undergo period shortening following exposure to a 50 µT MF. (e). 2-way ANOVA of *cry^M^* flies at 50 µT (3Hz) shows no significant MF based interaction F_1_,_180_=0.3, p=0.6).

Extended Data Figure 7

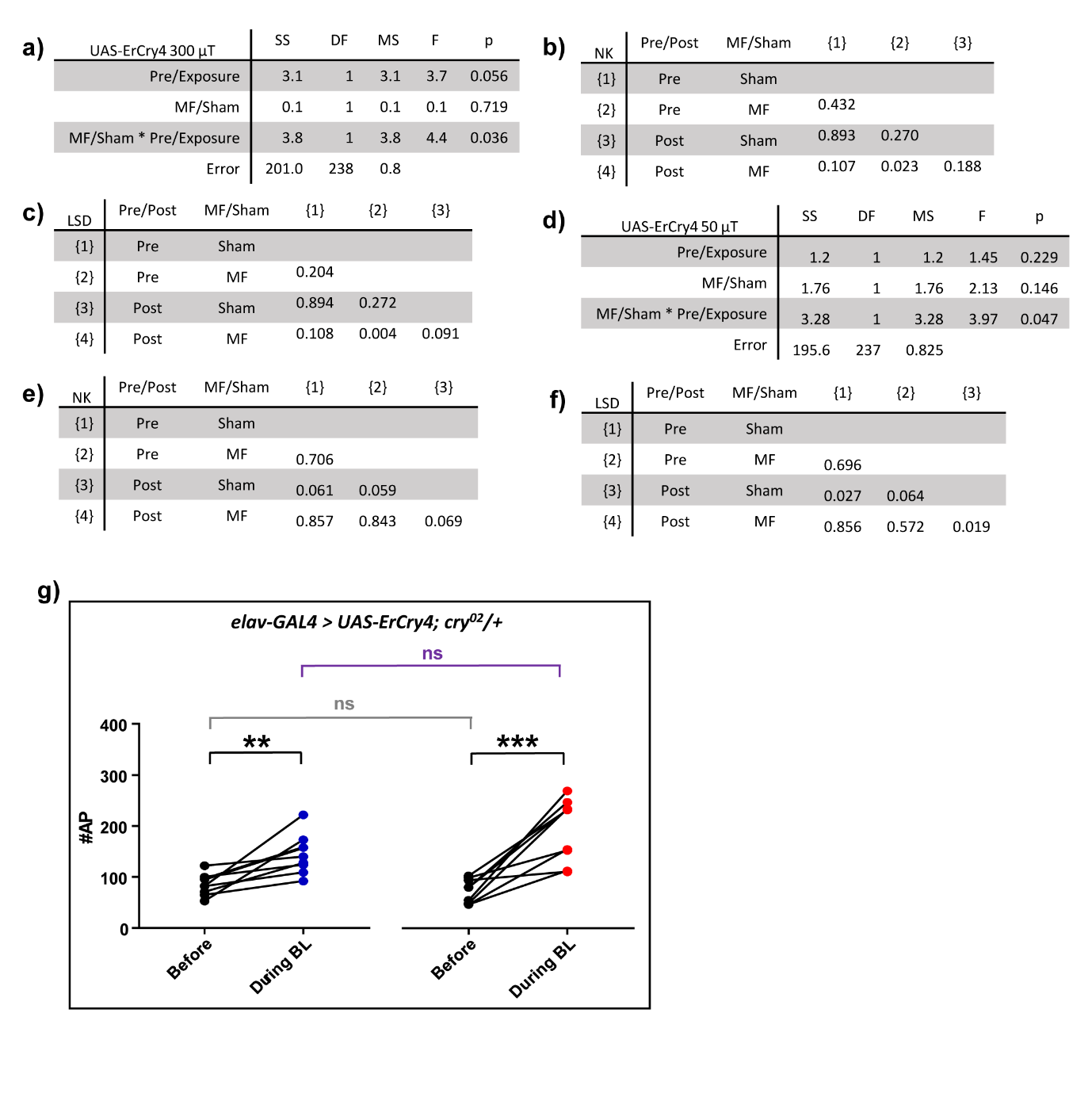

**Extended Data Figure 7.** Circadian and electrophysiological data for ErCry4 in main text Fig 4.

(a). 2-way ANOVA reveals a significant Sham/MF x pre/post-exposure interaction and a period shortening following exposure to a 300 µT / 3Hz MF. (b). A Newman-Keuls and (c). Fisher LSD post-hoc tests for ErCry4 at 300 µT / 3Hz MF. (d). A 2-way ANOVA also reveals a significant MF effect at 50 µT / 3Hz on ErCry4. (e). Newman-Keuls and (f). Fisher LSD post-hoc tests for ErCry4 at 50 µT / 3Hz MF exposure. (g). Raw AP counts for BL and BL+MF exposure for ErCry4, showing an increase in neuronal excitability to both stimuli. Paired t-tests were used to compare before *vs*. during for cells exposed to BL (left hand graph) or to BL±MF exposure (right hand graph). MF-potentiation between the two groups was tested by unpaired t-tests (different cells). ns p=>0.05, * p = ≤0.05, ** p=≤0.01, *** p=≤0.001.
